# CbpD crystal structure adds intrigue to substrate-specificity motifs in chitin-active lytic polysaccharide monooxygenases

**DOI:** 10.1101/2022.04.15.488525

**Authors:** Christopher M. Dade, Badreddine Douzi, Cristian Cambillau, Genevieve Ball, Romé Voulhoux, Katrina T. Forest

## Abstract

*Pseudomonas aeruginosa* secretes diverse proteins via its Type 2 Secretion System, including a 39 KDa Chitin-Binding Protein, CbpD. CbpD was recently shown to be a lytic polysaccharide monooxygenase active on chitin, and to contribute substantially to virulence. To-date no structure of this virulence factor has been reported. Its first two domains are homologous to those found in the crystal structure of *Vibrio cholerae* GbpA, while the third domain is homologous to the NMR structure of the *Cellvibrio japonicus Cj*LPMO10A CBM73 domain. We report the 3.0 Å resolution crystal structure of CbpD solved by molecular replacement, which required *ab initio* models of each CbpD domain generated by the artificial intelligence deep learning structure prediction algorithm RoseTTAFold. The structure of CbpD confirms previously postulated chitin-specific motifs in the AA10 domain while challenging the deterministic effects of other postulated substrate specificity motifs. Additionally, the structure of CbpD shows that post translational modifications occur on the chitin binding surface. Moreover, the structure raises interesting possibilities about how Type 2 Secretion System substrates may interact with the secretion machinery and demonstrates the utility of new artificial intelligence protein structure prediction algorithms in making challenging structural targets tractable.

## 1. Introduction

*Pseudomonas aeruginosa* secretes diverse proteins via its Type 2 Secretion System, including a 39 KDa Chitin-Binding Protein (CbpD) (Bleves *et al*., 2010). CbpD was originally identified by its ability to bind only crystalline chitin (Folders *et al*., 2000), and was thus classified in the Carbohydrate Binding Module 33 family (CBM33) (Cantarel *et al*., 2009). CbpD is a virulence factor (Askarian *et al*., 2021) that carries several different post-translational modifications (Gaviard *et al*., 2019; Ouidir *et al*., 2014; Gaviard *et al*., 2018) and whose function may be modulated by proteolytic cleavage after secretion through the Type 2 Secretion System (T2SS) (Park & Galloway, 1995; Braun *et al*., 1998; Folders *et al*., 2000).

A decade after the discovery of CbpD, it was shown that members of the CBM33 family are mono-copper binding enzymes with activity on recalcitrant polysaccharide substrates (Vaaje-Kolstad *et al*., 2010). These enzymes were reclassified as lytic polysaccharide monooxygenases (LPMOs) (Horn *et al*., 2012). Specifically, LPMOs catalyze the oxidation of β-1,4 glycosidic bonds in abundant natural polymers, including chitin, xylan, cellulose, and starch (Hamre *et al*., 2015; Forsberg *et al*., 2011; Vu, Beeson, Span *et al*., 2014; Corrêa *et al*., 2019), with a range of substrate selectivity (Zhou *et al*., 2019; Kojima *et al*., 2016; Agger *et al*., 2014; Isaksen *et al*., 2014) and regioselectivity (Vaaje-Kolstad *et al*., 2017; Li *et al*., 2012; Isaksen *et al*., 2014; Vu, Beeson, Phillips *et al*., 2014; Forsberg, Røhr *et al*., 2014). The decomposition of unstable oxidation intermediates results in glycosidic bond cleavage, nicking the polysaccharide and creating new ends for processive degradation of the biopolymer (Agostoni *et al*., 2017). While there has been some disagreement about the precise mechanism of LPMO oxidative cleavage (Bissaro *et al*., 2017; Hangasky *et al*., 2018), a thorough model has recently been proposed for the utilization of H_2_O_2_ as the reactive co-substrate (Bissaro *et al*., 2020). LPMOs are further characterized by a solvent exposed Peisach–Blumberg Type-2 copper active site in which a mononuclear copper ion is coordinated in a T-shape by a histidine brace comprising the N-terminal amine and two histidines (Peisach & Blumberg, 1974; Hemsworth, Taylor *et al*., 2013; Forsberg, Røhr *et al*., 2014).

The discovery of catalytic activity in CBM33 precipitated a reorganization of the Carbohydrate Active enZYme (CAZy) database and the creation of a new enzyme class—the auxiliary activity (AA) enzymes—into which CBM33 members and other enzymes were sorted (Levasseur *et al*., 2013). Based upon their sequence similarity, all CBM33 enzymes, including CbpD, were reclassified into the Auxiliary Activity 10 (AA10) family (Levasseur *et al*., 2013). LPMOs have now been found in bacterial, plant, fungal, archaeal, insect, and viral genomes (Beeson *et al*., 2015; Shukla *et al*., 2016; Beeson *et al*., 2012; Sabbadin *et al*., 2018) and are classified into families AA9–11 and AA13–16 (Vandhana *et al*., 2022).

A majority of LPMOs comprise a single copper-binding catalytic AA domain, while others have additional domains, themselves classified into families based on sequence identity (Vaaje-Kolstad *et al*., 2017; Horn *et al*., 2012; Book *et al*., 2014; Agostoni *et al*., 2017). These additional domains often contain aromatic residues that contribute to substrate binding and specificity (Gilbert *et al*., 2013; Cuskin *et al*., 2012; Crouch *et al*., 2016) and likely also protect LPMOs from autocatalytic inactivation (Courtade *et al*., 2018; Forsberg *et al*., 2018). In addition to its AA10 domain, CbpD has a GbpA2 domain and a Carbohydrate Binding Module 73 (CBM73), as annotated in PFAM (Mistry *et al*., 2021) and CAZy databases (Drula *et al*., 2022), respectively (Figure 1).

**Figure 1:**
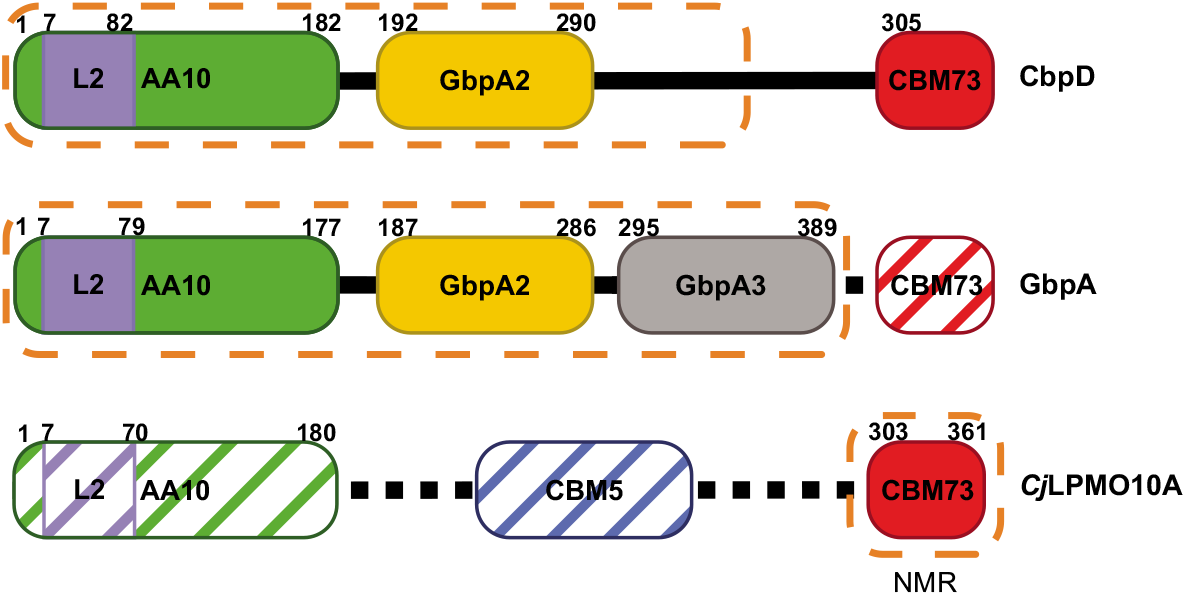
The domain architecture of CbpD and representative domain structures. CbpD is composed of 3 domains: an AA10 (green), a GbpA2 (yellow), and a CBM73 (red) domain. The domain structures of the AA10 and GbpA2 families are represented in the crystal structure of *V. cholera* GbpA (PDB:2XWX), while the domain structure of the CBM73 family is represented in the NMR solution structure of *Cj*LPMO10A (PDB:6Z40). Hashes represent regions not included in constructs used for cited structural studies. Dashed orange encapsulates the domain(s) resolved in the respective reported structures. Residue boundaries for each domain are labelled based on previous annotations, adjusted so the N-terminal His in each mature protein is residue 1.

While the catalytic activity of LPMOs is driving technological advances in biomass degradation (Harris *et al*., 2010; Arfi *et al*., 2014; Horn *et al*., 2012; Rani Singhania *et al*., 2021), their role in virulence has been less thoroughly investigated. LPMOs play a role in virulence (Wong *et al*., 2012; Loose *et al*., 2014; Chiu *et al*., 2015; Sabbadin *et al*., 2021; Agostoni *et al*., 2017) and are upregulated in pathogenic bacteria when exposed to human tissues (Vebø *et al*., 2009, 2010). Similarly, evidence has existed for two decades that CbpD might play a role in *P. aeruginosa* virulence. The CbpD gene has been found extensively in pathogenic clinical isolates (Folders *et al*., 2000) and is upregulated in the lungs of cystic fibrosis patients (Sriramulu *et al*., 2005). Recently, Askarian *et al*. demonstrated that catalytically active CbpD promotes *P. aeruginosa* survival in human whole blood, extends the survival of some other bacteria that do not endogenously express an LPMO, prevents cell lysis by the terminal complement pathway, and likely aids *P. aeruginosa* resistance to immune clearing in mouse infection models (Askarian *et al*., 2021).

These insights into the role LPMOs play in virulence have prompted studies of the structure and function of chitin-binding and virulence-associated LPMOs in pathogenic bacteria. The AA10 and GbpA2 domains of CbpD are represented structurally in the *Vibrio cholerae* colonization factor GbpA (PDB:2XWX) (Wong *et al*., 2012), while the CBM73 domain is found in a recently reported standalone NMR solution structure of the CBM73 domain from *Cellvibrio japonicus Cj*LPMO10A (PDB: 6Z40) (Madland *et al*., 2021). Despite availability of these structures of representatives of the three domains of CbpD, no structure of this virulence factor has been reported.

Concurrently with progress in understanding the role of LPMOs in virulence, recent advances in artificial intelligence (AI)-powered *ab initio* protein structure prediction (Baek *et al*., 2021*a*) allowed us to solve the partial structure of the virulence factor CbpD. While the structures of other LPMOs have been reported, many are single-domain LPMOs or are the catalytic domain of multidomain LPMOs (Vaaje-Kolstad, Houston *et al*., 2005; Vaaje-Kolstad *et al*., 2012; Gudmundsson *et al*., 2014; Forsberg, Røhr *et al*., 2014; Mekasha *et al*., 2016; Hemsworth, Taylor *et al*., 2013; Aachmann *et al*., 2012). While the structure of *V. cholerae* GbpA is multimodular, it was generated from a truncated construct lacking the C-terminal domain (a CBM73) (Wong *et al*., 2012) (Figure 1). We report the AI-enabled crystal structure determination of a multidomain LPMO. This structure of CbpD provides insights into domain architecture, substrate binding motifs, crystal packing, and apparent order and disorder propensities in multidomain LPMOs.

## 2. Methods and materials

### 2.1. Protein production and crystallization

CbpD from *P. aeruginosa* was previously cloned into the EcoR1 site of pT7.5 with an encoded C-terminal 10x His tag to yield pCbpD (Cadoret *et al*., 2014). Our purification procedure was modified only slightly from that publication. Overnight cultures of BL21 DE3 carrying pCbpD were used to inoculate 2 x 2L cultures of TB medium, which were incubated at 37°C at 220 rpm until OD_600_ reached 0.5-0.6. Protein expression was induced with 1mM IPTG, and proceeded overnight at 30°C. Cells were collected using low spin centrifugation, and the pellet was resuspended in buffer A (50mM Tris, 300mM NaCl, 10mM imidazole, pH 8.2) and frozen at −80°C. After 2 hours, cells were thawed in a 37°C water bath for 20–30 minutes to induce lysis. DNAse, MgSO_4_, and lysozyme were added to final concentrations of 20 μg/ml, 20mM, and 0.5 μg/ml, respectively. The mixture was stirred at 4°C for 1 hour to complete lysis. Clarified lysate was loaded onto a 5ml FFcrude prepacked nickel column (GE Healthcare) pre-equilibrated with buffer A. The column was washed with 10% buffer B (50mM Tris, 300mM NaCl, 250mM imidazole, pH 8.2), and CbpD was eluted using a gradient from 10–50% buffer B over 20 minutes. The elution fraction containing CbpD was concentrated using a 10K Amicon concentrator to 5 ml and loaded onto a Superdex S200 26/60 column equilibrated with 20mM Tris, 100mM NaCl pH 8. Fractions were analyzed by SDS-PAGE, and fractions containing CbpD were collected, pooled, and concentrated to 8 mg/ml (Supplementary Figure 1).

Crystals were grown by the hanging drop method against a reservoir solution of 0.2 mM di-ammonium tartrate in 20% (w/v) PEG 3350 (condition H2 from a Quiagen PEG sparse matrix screen). A crystal was harvested, mounted, and flash-cooled in liquid nitrogen without additional cryoprotection.

### 2.2. Data collection and structure determination

An x-ray diffraction dataset was collected to 2.98 Å resolution from a single crystal on the PROXIMA-1 beamline at the SOLEIL Synchrotron. The data set was processed in XSCALE (Kabsch, 2010). Initial attempts to solve the structure by molecular replacement with Phaser (McCoy *et al*., 2007) in Phenix (Liebschner *et al*., 2019), using the first two domains of the *V. cholerae* multidomain LPMO GbpA individually (PDB: 2XWX) (Wong *et al*., 2012) provided strong rotation function solutions, but no significant translation function outputs. The amino acid sequence of mature CbpD was then submitted to the Robetta, Phyre2 (Kelley *et al*., 2015), and PSIPRED (Buchan & Jones, 2019) servers, and 8 *ab initio* models were generated: 5 from RoseTTAFold (Baek *et al*., 2021*a*), 1 from Phyre2, and 2 from DMPFold (Greener *et al*., 2019). MR was attempted using each CbpD model as a template. While no full-length model was able to generate a high confidence solution, we chose the model with the highest LLG and TFZ scores (47 and 9 respectively) to break into domains. The AA10 domain of this RoseTTAFold model provided a partial solution (LLG: 181 TFZ: 18), which was held fixed in a subsequent round of MR using the RoseTTAFold GbpA2 output as the search model. This generated a successful MR solution (LLG: 432 TFZ: 22) containing the AA10 and GbpA2 domains of CbpD. An ensemble of CBM73 domains — including models generated by RoseTTAFold, Raptor-X (Askarian *et al*., 2021), and the NMR solution structure of the *Cellvibrio japonicus* CBM73 (Madland *et al*., 2021) — was then used in subsequent rounds of MR, holding fixed the AA10/GbpA2 partial solution. Despite numerous attempts using varied input parameters, only low-likelihood solutions were ever found for the CBM73 and upon visual inspection were unlikely due to clashes with either the partial model of CbpD or symmetry mates.

The two-domain model of CbpD was rebuilt using Phenix.autobuild (Liebschner *et al*., 2019) to create the linker region between the AA10 and GbpA2 domains. The electron density was good enough to manually extend the model past the GbpA2 domain to residue 296 in Coot (Emsley *et al*., 2010), which was used with Phenix.refine (Liebschner *et al*., 2019) for iterative rounds of real and reciprocal-space refinement. Refinements were performed with a high-resolution cutoff of 3 Å. Atomic displacement parameters were refined as group factors until the final round of refinement, when they were refined using one translation-libration-screw (TLS) group (Urzhumtsev *et al*., 2013). The final model refined to R_cryst_ and R_free_ factors of 0.21 and 0.25, respectively (Table 1).

**Table 1.**
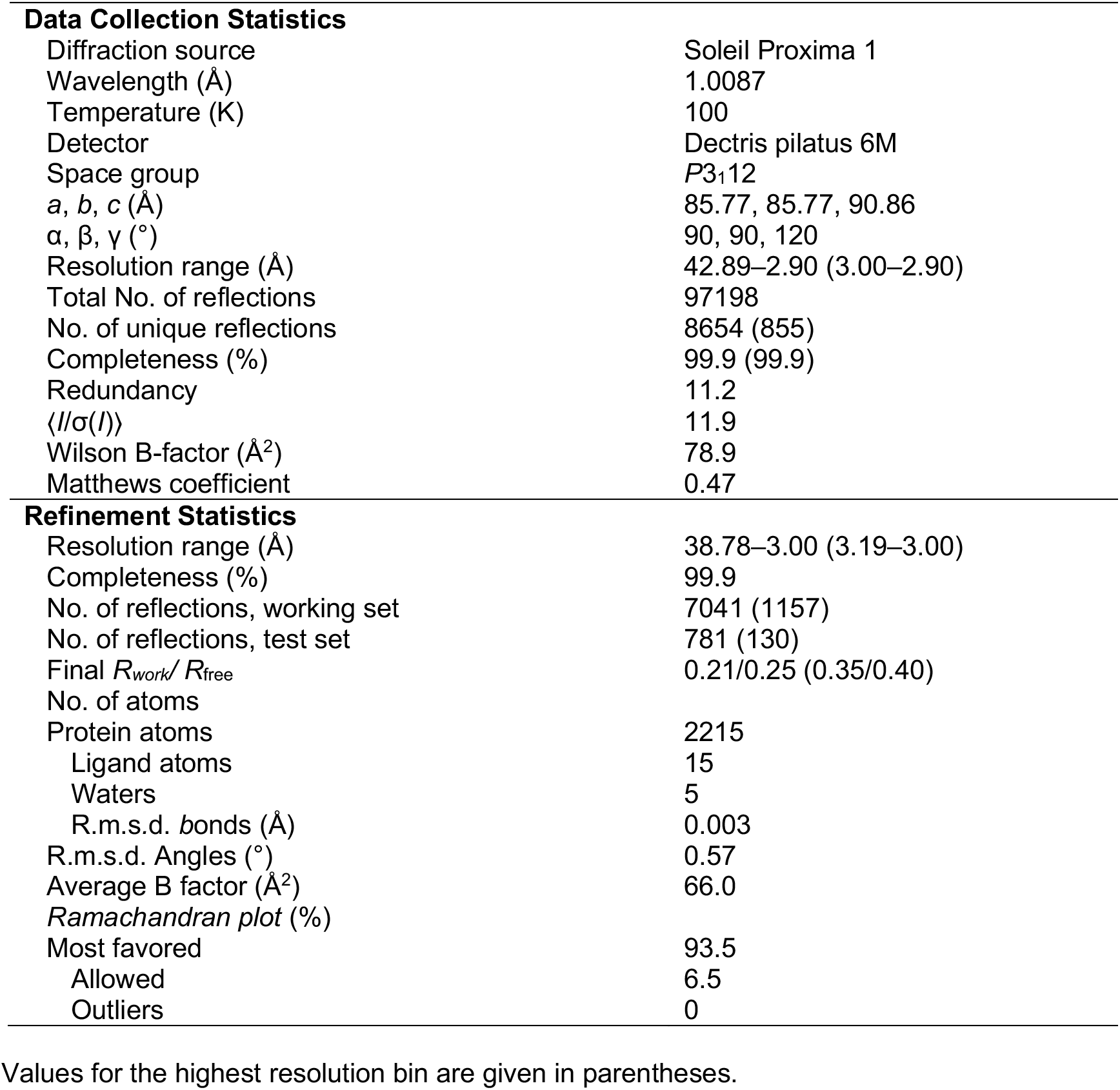
Data collection, processing, and refinement

|  |  |
| --- | --- |
| **Data Collection Statistics** |  |
| Diffraction source | Soleil Proxima 1 |
| Wavelength (Å) | 1.0087 |
| Temperature (K) | 100 |
| Detector | Dectris pilatus 6M |
| Space group | $P3_112$ |
| $a, b, c$ (Å) | 85.77, 85.77, 90.86 |
| $\alpha, \beta, \gamma$ (°) | 90, 90, 120 |
| Resolution range (Å) | 42.89–2.90 (3.00–2.90) |
| Total No. of reflections | 97198 |
| No. of unique reflections | 8654 (855) |
| Completeness (%) | 99.9 (99.9) |
| Redundancy | 11.2 |
| $\langle I/\sigma(I) \rangle$ | 11.9 |
| Wilson B-factor (Å<sup>2</sup>) | 78.9 |
| Matthews coefficient | 0.47 |
| **Refinement Statistics** |  |
| Resolution range (Å) | 38.78–3.00 (3.19–3.00) |
| Completeness (%) | 99.9 |
| No. of reflections, working set | 7041 (1157) |
| No. of reflections, test set | 781 (130) |
| Final $R_{\text{work}}/R_{\text{free}}$ | 0.21/0.25 (0.35/0.40) |
| No. of atoms |  |
| Protein atoms | 2215 |
| Ligand atoms | 15 |
| Waters | 5 |
| R.m.s.d. bonds (Å) | 0.003 |
| R.m.s.d. Angles (°) | 0.57 |
| Average B factor (Å<sup>2</sup>) | 66.0 |
| Ramachandran plot (%) |  |
| Most favored | 93.5 |
| Allowed | 6.5 |
| Outliers | 0 |

Values for the highest resolution bin are given in parentheses.

### 2.3. Modeling and Docking

The orientation of CbpD on chitin could be confidently predicted because all chitin-active LPMOAA10s cleave at C1—thus the reducing end of chitin must align with E171, and the non-reducing end must align with Y40/E20 (see Bissaro *et al*. 2018 for an in-depth review of LPMO hydrolysis products, regioselectivity, and substrate orientation).

In order to evaluate the possibility for packing of the CBM73 domain in the CbpD crystal lattice, the RoseTTAFold model of the CbpD CBM73 domain (Baek *et al*., 2021*a*) was first manually docked into the remaining space using Coot (Emsley *et al*., 2010) to minimize clashes with symmetry mates. A surface model blob of each CBM73 domain was then generated in Pymol (Schroöinger, 2015) by setting the B-factor for all atoms to 100, setting the gaussian resolution to 8.0, generating a gaussian map for each CBM73 domain (grid=1, buffer=6), and finally generating an isosurface for each CBM73 domain (surface quality=1).

To assess the fit of structural models to experimental small angle scattering envelopes, a 15 Å volume map was generated in ChimeraX (Pettersen *et al*., 2021) by converting the envelope bead model of CbpD (Askarian *et al*., 2021) (SASBDB ID: SASDK42) into a density map using the molmap command. CbpD models were then manually placed into the density map and their position refined in the density map using the “Fit in Map” tool in ChimeraX.

3D structure figures were generated in Pymol (Schrödinger, 2015). Structurebased topology was depicted through use of Pro-origami (Stivala *et al*., 2011*a*).

## 3. Results

Models generated from newly introduced protein structure prediction AI algorithms enabled us to solve the structure of CbpD using molecular replacement, which had not been successful using the representative domain structures shown in Figure 1. The structure of mature CbpD residues 1–296 (where 1 is the N-terminal His after signal peptide cleavage) was determined to 3.0 Å resolution with a single molecule in the asymmetric unit (Table 1). The overall structure contains the AA10 and GbpA2 domains, a well-ordered linker between them, and a well-ordered portion of the linker between the GbpA2 and CBM73 domains out to residue 296 (Figures 2A and 2B). Residues 297–374 (the remainder of the linker and the CBM73 domain) were not visible in the electron density map. We compared the AI-predicted structure to the final CbpD structure and found an all-atom RMSD of 11.3 Å (2215 atoms). We then compared each domain and found all-atom RMSDs of 1.5 Å (1370 atoms) and 1.6 Å (733 atoms) for the AA10 and GbpA2 domains, respectively (Figure 2C), compared to 8.9 Å (874 atoms) and 4.6 Å (569 atoms) for refined CbpD domains and the representative domains in GbpA. In hindsight, it is noteworthy that a fern insecticidal AA10 (Yadav et al., 2019; PDB code 6IF7) has very low RMSD of 1.5 Å over 1252 atoms compared to the CbpD AA10, and of those AA10s whose structures are now available, is the most close in sequence to CbpD (Figure S2). This structure was not yet available when our x-ray data were collected but might have proved useful in phase determination by molecular replacement.

**Figure 2.**
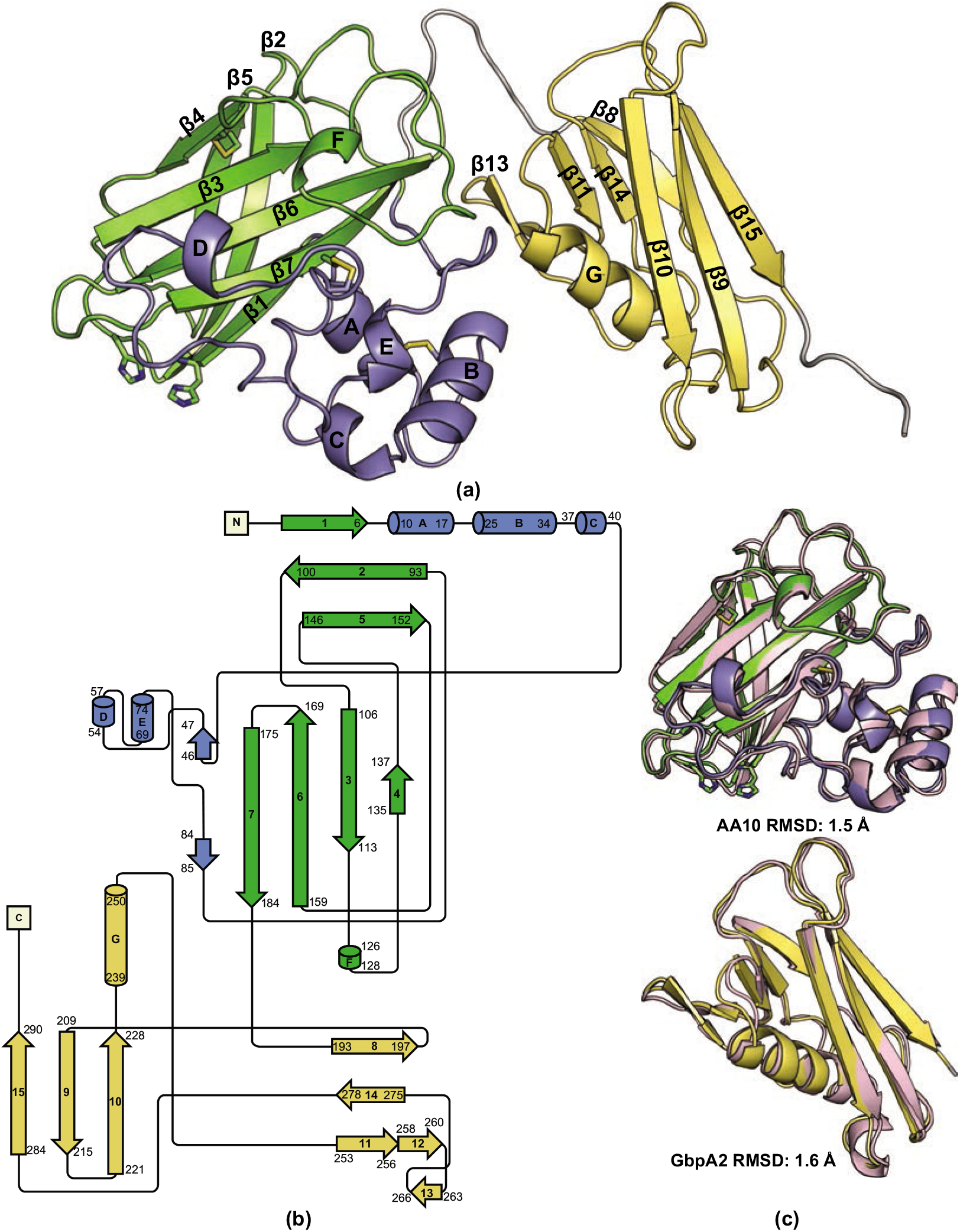
The overall structure of CbpD. A) The structure of CbpD contains an AA10 domain (green) with L2 region (slate) between β_1_ and β_2_ connected to a GbpA2 domain (yellow) by flexible linker regions (grey). Disulfide bridges and the active site His residues are shown as sticks. B) The structure-based topology diagram for CbpD was initially generated with Pro-origami (Stivala *et al*., 2011*b*), and long intervening stretches were shortened manually and are not to scale. C) A comparison of the AA10 and GbpA domains from CbpD structure and predicted structural domain models from RoseTTAFold (Baek *et al*., 2021*b*) (pink) reveals a high degree of predictive accuracy. The all-atom RMSDs between the predicted and crystallographic AA10 and GbpA2 domains are 1.5 Å and 1.6 Å, respectively.

### 3.1 CbpD AA10 Domain

The AA10 domain contains a typical LPMO AA10 fold, with a central 7-stranded β-sandwich comprising two anti-parallel β-sheets connected by loop regions and an L2 subdomain between β_1_ and β_2_ (Figures 2A,B). This L2 region forms the majority of the substrate-binding surface and displays a majority of the sequence variation in LPMOAA10s (Wu *et al*., 2013; Forsberg *et al*., 2016; Bissaro *et al*., 2017; Vaaje-Kolstad *et al*., 2017). The AA10 domain also contains three disulfide bonds: one linking β4 to β5 (Cys135 and Cys151), one linking two helices in the L2 domain (Cys14 and Cys27), and one linking the L2 domain to the β-sandwich through β7 (Cys64 and Cys179).

#### Copper-binding active site primary coordination sphere

The active site comprises the canonical histidine brace found in LPMOs and is formed by His1 and His104 (Figure 3A). Although the active site did not appear to contain a coordinated metal, a copper could be modeled into the active site with the electron density contoured to 0.7 σ (Figure 3A). Missing or partially occupied active site Cu is common in LPMOAA10 structures (Frandsen & Lo Leggio, 2016). In this position, the copper would only be coordinated axially in a T-shape by the histidine brace, as expected in an LPMOAA10 (Forsberg, Røhr *et al*., 2014). The primary coordination sphere (Figure 3B) also includes highly conserved features of LPMOAA10s that help structure the active site. This includes a phenylalanine (Phe176) axial to the active site and a universally conserved alanine (Ala102) (Figure 3C) (Hemsworth, Davies *et al*., 2013; Forsberg, Røhr *et al*., 2014) that is believed to restrict coordination of cosubstrates in the solvent-exposed axial position and increase the likelihood an oxygen cosubstrate coordinated at the equatorial position will be activated for catalysis, thus conferring C1/C4 oxidizing specificity (Forsberg, Mackenzie *et al*., 2014; Borisova *et al*., 2015). A well conserved tryptophan (Trp167) among LPMOAA10s (Forsberg, Røhr *et al*., 2014; Meier *et al*., 2018) thought to protect the active site against oxidative inactivation during uncoupled turnovers (Loose *et al*., 2018; Paradisi *et al*., 2019; Gray & Winkler, 2021) is also present (Figures 3B,C).

**Figure 3.**
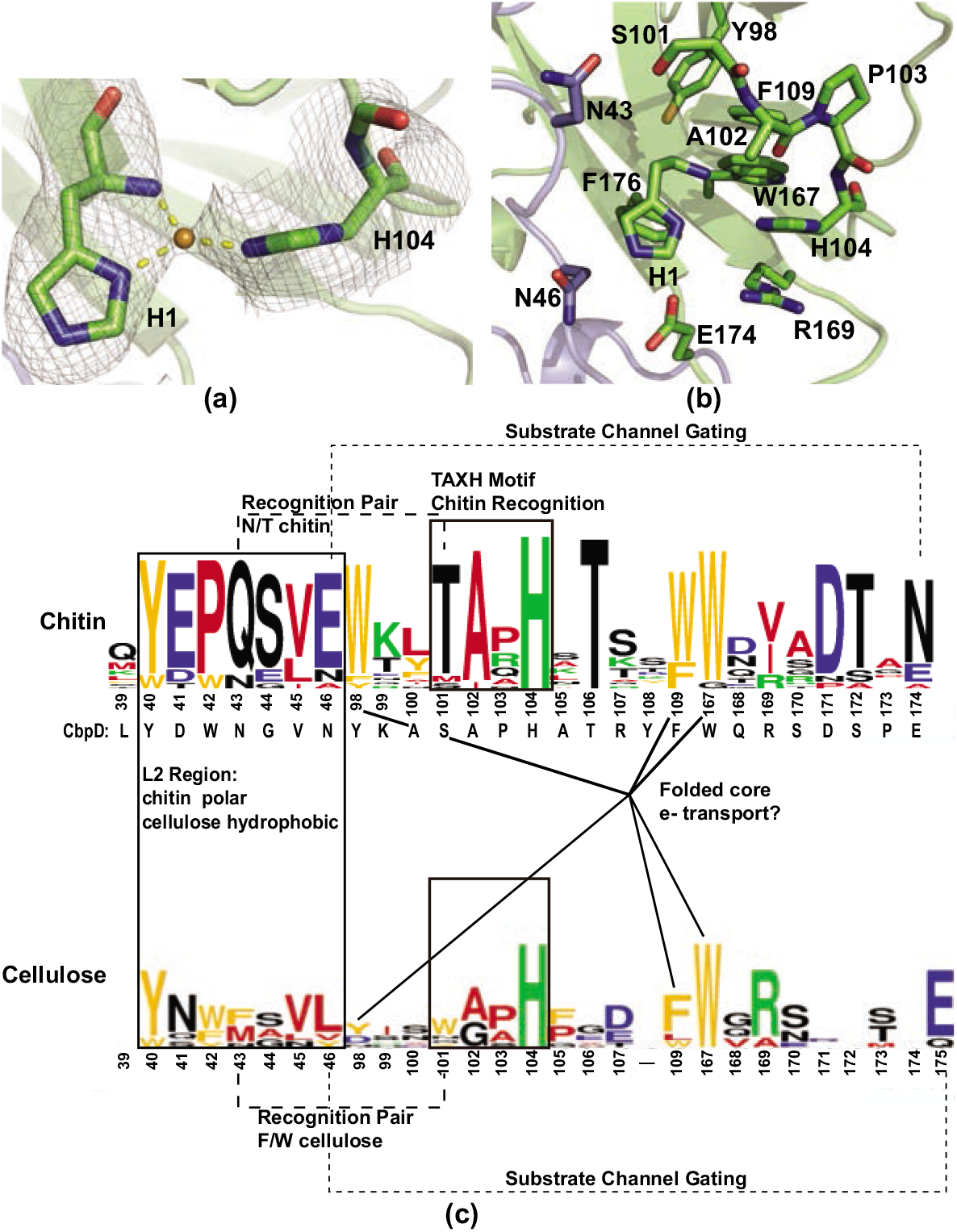
The structure of CbpD reveals some chitin-specific motifs are preserved while others are more similar to cellulose-specific motifs. A) The active site of CbpD contains a canonical His brace motif. 2Fo-Fc electron density contoured at 0.7σ allows Cu to be modelled into the active site, approximately 2 Å away from the three coordinating N, as expected. B) The CbpD active site also contains a number of highly conserved residues. C) Motifs previously identified as conserved among and/or proposed to confer substrate specificity in chitin- and cellulose-active LPMOAA10s are mapped to sequence logos generated from multiple sequence alignments of 13 chitin- and 7 cellulose-active LPMOAA10s listed as experimentally validated in the CAZy database (Drula *et al*., 2022) and in Zhou and Zhu 2020, with the addition of CbpD to the chitin-active sequences. Motifs were concatenated for display, and residues are numbered as in CbpD for the chitin-active sequence logo. In the cellulose-active sequence logo, residues are number by the structurally equivalent position in CbpD. The CbpD amino acid identities are below the chitin Logo. MSAs were generated with the T-coffee web server (Notredame *et al*., 2000; di Tommaso *et al*., 2011). Sequence logos were generated with WebLogo (Crooks *et al*., 2004).

The active site of CbpD, however, deviates from or lacks two features previously postulated to be conserved among chitin-active LPMOAA10s. CbpD lacks the cluster of 3 tryptophans previously identified as conserved among chitin-active AA10s. These are Tyr98, Phe109, Trp167 in CbpD (Figure 3C). This tryptophan cluster was thought to be critical in a catalytic mechanism model based on O_2_ as the oxygen cosubstrate that required external electron-transfer (ET) (Forsberg, Røhr *et al*., 2014). An internal electron tunneling path was putatively identified in fungal LPMOAA9s and was thought to connect the active site to either a surface exposed lysine through a hydrogen bond network or a surface patch through aromatic residues where Cellobiose Dehydrogenase (CDH) could bind as an external electron donor (Li *et al*., 2012). An equivalent ET pathway has never been identified in bacterial chitin-active LPMOs (Vaaje-Kolstad *et al*., 2012), however, and a number of alternative electron donor sources have now been identified for LPMOs, including small molecules (Askarian *et al*., 2021; Westereng *et al*., 2015, 2016), photo-active pigments (Cannella *et al*., 2016), and even substrates themselves (Westereng *et al*., 2015). Subsequent experimental (Bissaro *et al*., 2017; Kuusk *et al*., 2018, 2019; Jones *et al*., 2020) and computational (Bissaro *et al*., 2020; Wang *et al*., 2018) studies have also challenged whether O_2_ is the natural cosubstrate in favor of H_2_O_2_. An H_2_O_2_-based mechanism would not require an external electron donor after the initial reduction of Cu from Cu(II) to Cu(I) before H_2_O_2_ binds (Bissaro *et al*., 2018*a*). Additionally, an ET pathway in LPMOAA10s would require electrons to transverse a conserved Phe (Phe176), which does not facilitate ET (Beeson *et al*., 2015). The conservation of this active site Phe and lack of conserved tryptophans cast further doubt that chitin-active LPMOAA10s rely on an internal ET pathway for catalysis. Interestingly a cellulose-active LPMOAA10 with this pattern of three aromatic residues has been identified (Forsberg, Røhr *et al*., 2014). This, combined with their location away from the putative substrate binding surface, indicates the Trp trio is also likely not a determinant in substrate specificity and may rather play a role in stabilizing the hydrophobic core of the domain.

#### Copper-binding active site secondary coordination sphere

The secondary shell of the CbpD active site also contains three highly conserved features of LPMOAA10s. First, In CbpD, the reportedly invariant TAXH chitin recognition motif (Forsberg *et al*., 2018), where H is the second histidine in the histidine brace (His104), is SAPH, with the conservative replacement of serine for threonine (Figures 3B,C). Second, CbpD fulfills the Gln–Thr pair previously identified as conserved among chitin-binding LPMOAA10s (Zhou & Zhu, 2020) with Asn43–Ser101 (Figures 3B,C). In cellulose-active AA10s, this pair is replaced by two large, hydrophobic residues, generally Phe–Trp (Forsberg *et al*., 2016). Third, previous molecular modeling of a chitin-active LPMOAA10 bound to crystalline chitin also identified a Glu–Asn pair (Glu60–Asn185 in *SmAA10A)* that appears to gate a substrate tunnel from the bulk solvent into the copper active site when an LPMOAA10 is bound to crystalline chitin (Figure 3C) (Bissaro *et al*., 2018*a*). This gated tunnel is proposed to be the route through which only small molecule cosubstrates (*e.g*., O_2_, O_2_•–, H_2_O_2_, or H2O) could access the active site. This Glu has been identified as conserved in all LPMOs as either a Glu or Gln and is required for enzymatic activity (Vaaje-Kolstad, Horn *et al*., 2005; Harris *et al*., 2010), likely by coordinating H_2_O_2_ and reactive oxygen species intermediates (Bacik *et al*., 2017; Bissaro *et al*., 2020; O’Dell *et al*., 2017; Span & Marletta, 2015; Hedegård & Ryde, 2017). CbpD contains an analogous Glu–Asn pair (Glu174–Asn46), but of note is that the two residues are flipped (Figures 3B,C), a feature also present in *Cj*LPMO10A (Forsberg *et al*., 2016). Nevertheless, the properties and position of the gating pair in the AA10 domain are retained, and despite being quite distant in the primary sequence from the putatively conserved position, Glu174 still points toward the active site of CbpD, approximately 5 Å away from the Cu site (Figures 3, 4A). This indicates Glu174/Asn46 may play a similar role in gating access to the active site when CbpD is bound to crystalline chitin. In these three chitin recognition motifs, side-chain properties are conserved.

**Figure 4.**
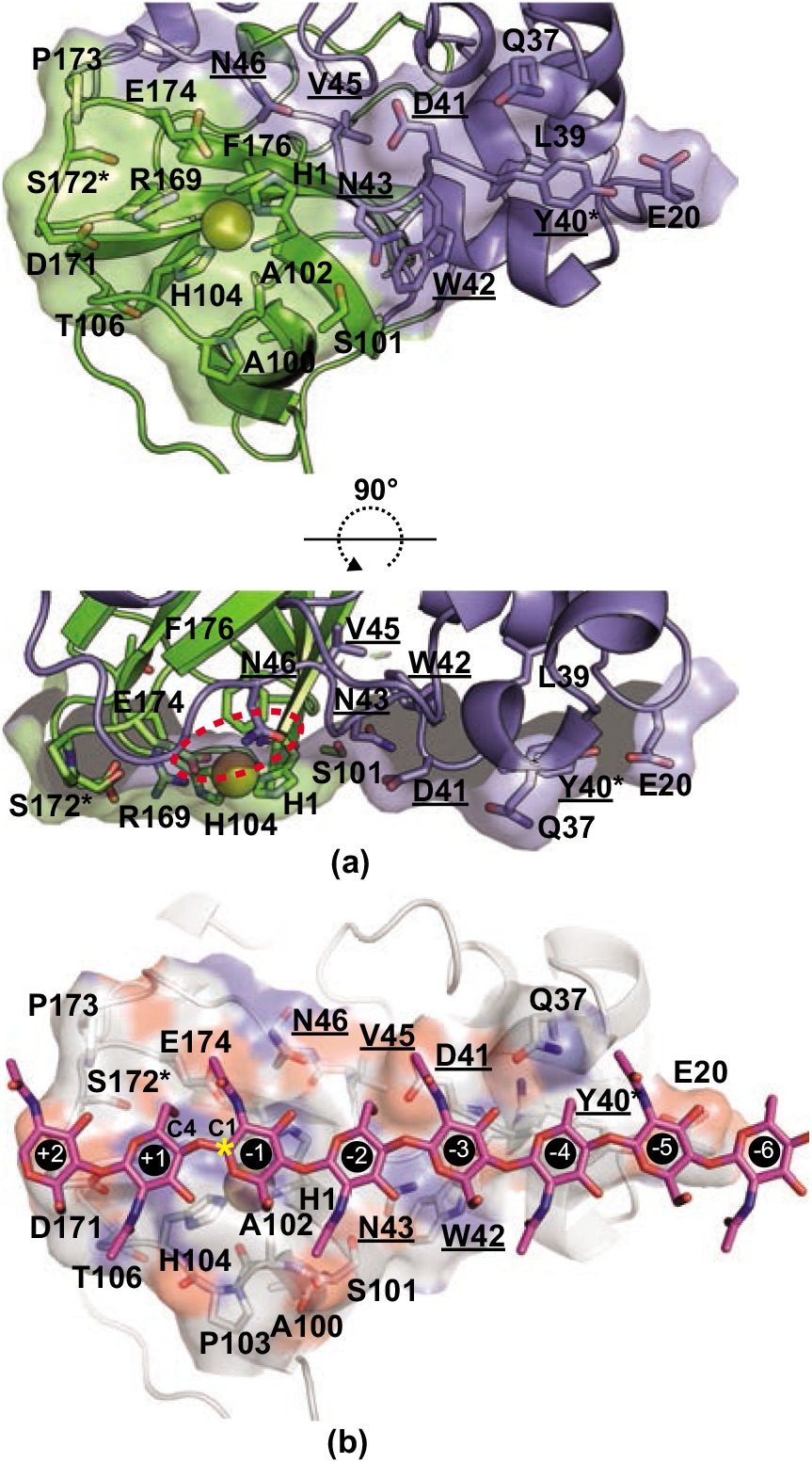
The chitin-binding surface of CbpD. A) Looking at the chitin-binding surface from the perspective of the crystalline substrate, residues identified in other LPMOs as interacting with chitin are labeled. Cu is modeled into the active site His brace for reference and orientation. Residues in the putative chitin-specific L2 motif are underlined. Ser172 and Tyr40, which are phosphorylated when CbpD is expressed and secreted by *P. aeruginosa*, are marked with an asterisk. B) Side view of the chitin-binding surface showing the relatively flat substrate-binding surface of the CbpD AA10 domain and the large role the L2 region (slate) plays in substrate binding. The substrate tunnel gating pair Asn46/Glu174 are indicated with a dashed oval. C) Based on the modeling of *Sm*AA10 interacting with β-chitin (Bissaro *et al*., 2018*b*), CbpD was manually docked onto crystalline anhydrous β-chitin (Nishiyama *et al*., 2011) (Crystallography Open Database: 1501776) Crystallography Open Database: 1501776) so that the substrate binding surface of CbpD (oxygen and nitrogen atoms of sidechains in red and blue respectively) was parallel to the chitin surface and aligned such that the chitin ran from one end of the surface (Glu171) across the active site to the opposite end (Tyr40/Glu20). A chito-octaose (NAG_8_) fragment interacting with subsites −6 to +2 is shown as sticks. Subsites are numbered following the standard practice for glycoside hydrolases (Sunna *et al*., 1997).

The secondary coordination sphere in CbpD also contains a reversed putative discriminating feature between chitin and cellulose-active LPMOAA10s. Chitin-active LPMOAA10s were identified to contain a short, aliphatic residue (generally an isoleucine or valine) just below the histidine brace, while that position was an Arg in cellulose-active LPMOAA10s (Figure 3) (Forsberg, Mackenzie *et al*., 2014; Forsberg, Røhr *et al*., 2014). It has previously been postulated that a short aliphatic residue in this position creates a cavity on the substrate binding surface of chitin-active LPMOAA10s that could accommodate a dioxygen cosubstrate for catalysis or the N-acetyl group found in chitin (Hemsworth, Taylor *et al*., 2013; Forsberg, Mackenzie *et al*., 2014). In chitin-active CbpD, however, this position is Arg169, and the cavity is further filled by Asp171 and Glu174 (Figure 4A). This is the case for chitin-active *Cj*LPMO10A as well (Forsberg *et al*., 2016). In fact, in 16.7% of reviewed chitin-active LPMOAA10s in the CAZy database (Drula *et al*., 2022), this position is an Arg (Figure S3). This raises questions about whether this residue is important for substrate discrimination and casts doubt on the importance of a surface cavity for catalytic activity.

#### Chitin-binding surface

The structure of CbpD reveals a canonical relatively flat surface of ~1000 Å^2^ encompassing the copper active site that is the presumed chitin-binding surface of CbpD (Figure 4A,B). While no studies have directly examined the role residues along this surface play in mediating chitin binding by CbpD, the structure of CbpD enables the mapping of homologous chitin-interacting residues from other characterized LPMOAA10s to CbpD. Residues that interact with chitin in chitin-specific LPMOAA10s have been identified through both HSQC NMR (Vaaje-Kolstad *et al*., 2010) and MD simulations (Bissaro *et al*., 2018*a*). The putative chitin-binding residues of CbpD could be identified by mapping these sets of residues onto CbpD (Tyr40, Asp41, His104 from both methods; Leu39, Asn43, Gly44, Ala100, Ser101, Ala102, Thr106 from NMR; and Asn46, Asp171, Glu174 from MD) (Figure 4A,B). These residues form a surface that extends from the conserved single aromatic residue (Tyr40) found on the substrate binding surface of chitin-active LPMOAA10s (Zhou & Zhu, 2020) to Asp171 at the edge of the substrate-binding surface. The structure suggests that Glu20 may further extend the substrate surface beyond Tyr40 in CbpD.

Interestingly, mapping features of chitin-specific LPMOAA10 substrate surfaces to CbpD revealed deviations in CbpD from previously identified trends. A motif on the L2 domain, beginning with the conserved substrate-binding Tyr and running down the center of the substrate-binding surface, purportedly exhibits differing polar characteristics in chitin- and cellulose-specific LPMOAA10s (Zhou *et al*., 2019). In chitin-specific LPMOAA10s examined, this motif contained at least 70% polar residues, with a consensus sequence of Y(W)EPQSVE (Figure 3C). Cellulose-specific LPMOAA10s examined had a motif containing more than 70% hydrophobic residues. In CbpD, this L2 motif is YDWNGVN beginning with the conserved Tyr40. This sequence differs from the consensus chitin-specific sequence, and the side chain properties fall between the chitin- and cellulose-specific motif properties (Figure 3C). This motif in CbpD, however, still forms a flat surface that appears to be an appropriate area for substrate binding (Figure 4B). The divergence of these residues in CbpD from previous trends raises questions about the extent to which this motif in LPMOAA10s determines substrate specificity. Interestingly, the chitin-active insecticidal putative LPMOAA10 produced by the fern *Tectaria macrodonta* (Tma12) has the sequence YEWNEVN closely matches the apparent outlier CbpD sequence here, providing further evidence that the Y(W)EPQSVE motif is not the determinant of chitin-specificity.

### 3.2 CbpD GbpA2 Domain

The second domain of CbpD is, as expected, similar to the GbpA2 domain structure GbpA (Wong *et al*., 2012) with an all-atom RMSD of 4.6 Å for 569 atoms (residues 193–292 in CbpD and 211–314 in GbpA). The GbpA2 domain comprises an α-helix partially sandwiched between two anti-parallel β-sheets (β-strands 9,10,15 and 8,11,14), with an unusual bend that allows β_11_ to pair with β_14_ while its immediate extension β_12_ interacts with the short β_13_ to form a hairpin above the helix (Figure 2A,B). The orientation of the GbpA2 domain relative to the AA10 domain is shifted in the structure of CbpD by 1 Å and 135.6° compared to the structure of GbpA (Figure 5A). This shift means the CbpD crystal structure is closer to the elongated conformation observed for both GbpA and CbpD by solution scattering (Wong *et al*., 2012; Askarian *et al*., 2021) (Figure S4). In contrast, GbpA adopts a U-shaped conformation in its crystal, forming an extensive interface in which each subunit contributes 3,120 Å^2^ to the dimerization interface, as calculated with PISA (Krissinel & Henrick, 2007). Notably, while CbpD does not form a similar dimer, it does display an intriguing continuation of the β-sheet from β15 of one CbpD GbpA2 domain into its symmetry mate, burying 990 Å^2^ per monomer (Figure 5B).

**Figure 5.**
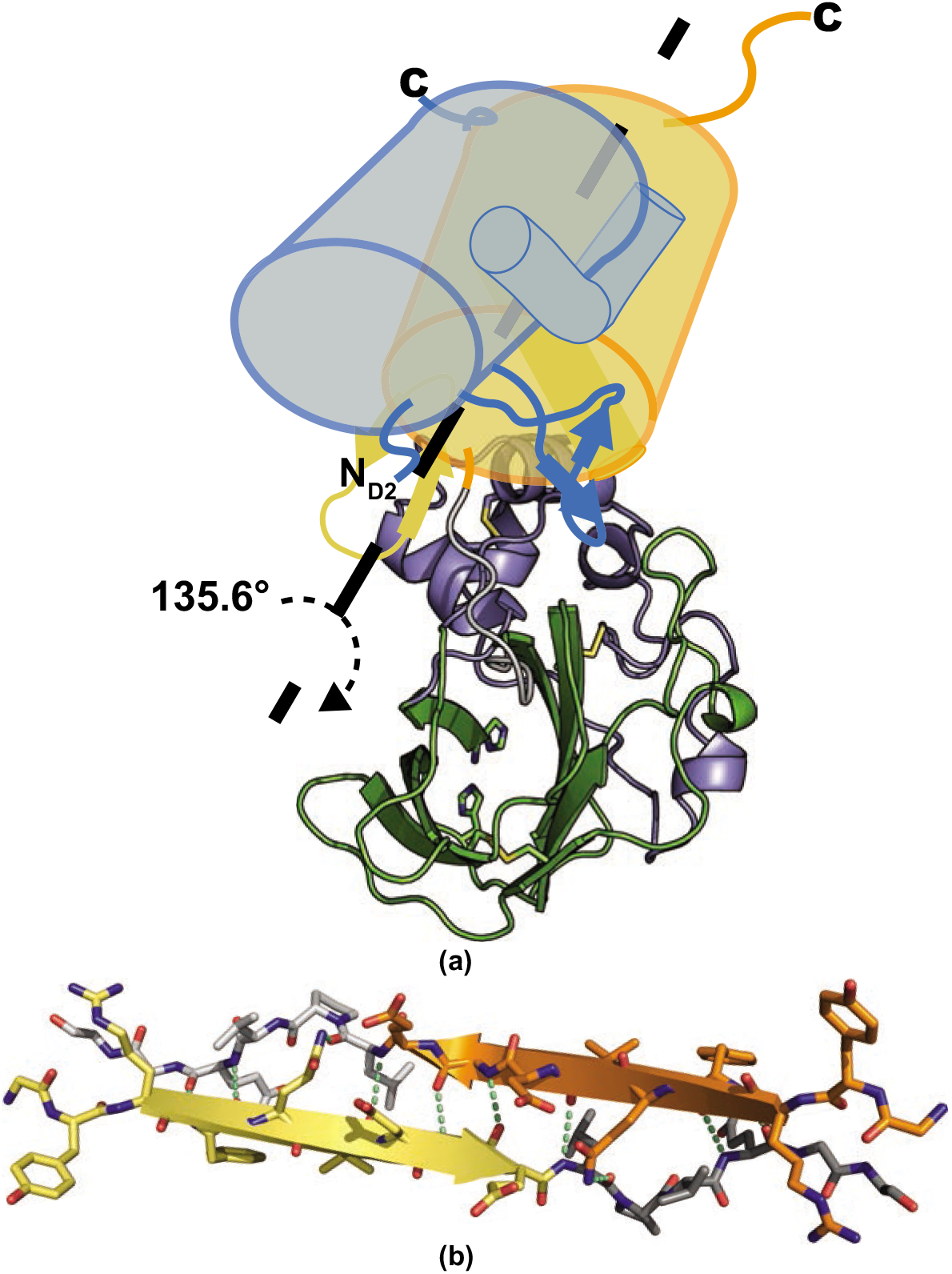
The role of the GbpA2 domain in crystal packing. A) The GbpA2 domain in the CbpD structure (yellow cylinder) is displaced by 1 Å and rotated by 135.6° relative to the GbpA2 domain in the GbpA structure (light blue cylinder) when their AA10 domains are aligned. The αG of each GbpA2 domain (smaller cylinders), the β_12_-β_13_ hairpins (stylized cartoons), and the C-termini are indicated for orientation. The AA10 domain of CbpD is represented as in Figure 2A. The displacement of GbpA2 domains is also noted by the ND2 label at the shifted N-terminus of this domain in GbpA. B) β_15_ and part of the linker between the GbpA2 and CBM73 domains of one CbpD (yellow and dark grey) engage in extensive β-strand interactions with a symmetry mate (orange and light grey) to form an extended β-sheet, burying 990 Å^2^ of surface area per monomer.

### 3.3 CbpD CBM73 Domain

While full-length CbpD-His_10_ was crystallized, good electron density was only observed through residue 296 (Figure 6A), leaving nine residues of the linker between the GbpA2 and CBM73 domains as well as the CBM73 domain itself unmodeled. Despite the absence of strong density to place the CBM73 domain, it is unlikely that the residues were missing from the protein crystal. Given the 10xHis affinity tag was attached to the C-terminus of CbpD, it is clear that full-length CbpD was purified via metal affinity chromatography (see Methods and Figure S1). Crystal solvent content analysis with Phenix.xtriage (Zwart *et al*., 2005) calculated 46% solvent content for a unit cell containing 374 residues, consistent with crystallization of full-length CbpD-His_10_. While there is some evidence CbpD is processed from its C-terminus by LasB into the 23kDa LasD when secreted by *P. aeruginosa* (Braun *et al*., 1998), CbpD-His_10_ was expressed in *E. coli*, which lacks LasB. The processing of CbpD into LasD would, moreover, result in the loss of all but the first 24 residues of the GbpA2 domain, which is fully present in the crystal structure. Additionally, there is a sufficiently large volume remaining in the crystal packing of CbpD to arrange the CBM73 domain (Figure 6B), although in a more compact configuration than the fully elongated configuration deduced from SAXS data (Askarian *et al*., 2021).

**Figure 6.**
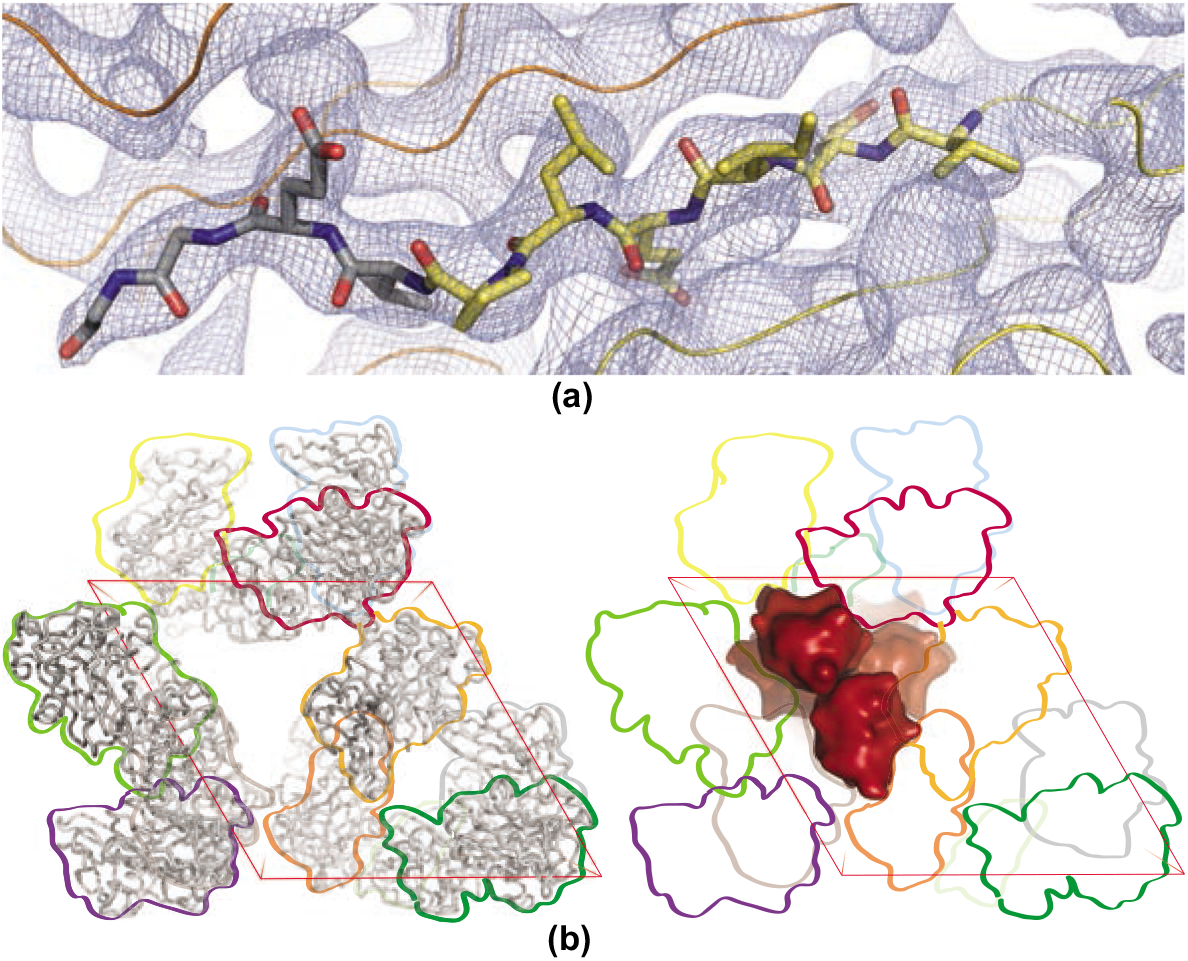
The CBM73 domain is present but unresolved in the CbpD crystal structure. A) The 2Fo-Fc electron density map is contoured at 1.0σ. The Cα trace of the CbpD GbpA2 domain is shown (yellow and grey with 287-296 as sticks) with symmetry mate (orange). The electron density extends to residue 296. B) Despite the absence of strong density to place the CBM73 domain, there is space in the crystal packing where the domain fits. Looking roughly down the c-axis of the unit cell, the left panel depicts one unit cell of the modeled residues of CbpD (ribbons) with each symmetry mate outlined for clarity and reference in the right panel (colors arbitrary). The right panel depicts how the CBM73 domain (red blobs) could pack into the remaining crystal volume.

## 4. Discussion

The structure of CbpD reported here is the first structurally determined LPMO from *P. aeruginosa* and highlights the utility new, AI-powered, *ab initio* protein structure prediction algorithms have in the structural biology toolkit. As demonstrated with CbpD, these *ab initio* models may help solve previously recalcitrant x-ray crystallography structures through MR by providing starting models that are substantially closer to the actual protein structure than homologous structures. Because these AI-generated models may provide better MR models, they may also enable protein structures to be solved with lower resolution or lower quality data than was previously routine. Additionally, these models will provide another path toward phasing protein structure targets without the need for additional experimentation steps such as Se-Met labeling, the collection of multiple datasets, or heavy metal soaking (which is also a danger to human and environmental health). These AI-generated models could be particularly beneficial for difficult crystal targets that are difficult to express and purify in large quantities, produce few high-quality diffracting crystals, or lack closely homologous structures that otherwise could serve as the basis for MR. The close agreement between AI-generated *ab initio* models and the crystal structures of each domain of CbpD, specifically the AA10 catalytic domain, also indicates that these AI results may provide high quality structural models for the thousands of LPMOs that have been identified but lack experimental structures to develop more robust hypotheses about the characteristics of LPMOs that affect catalysis and substrate selectivity.

The structure of CbpD also provides greater insight into the conserved features of chitin-active LPMOAA10s. CbpD displays a number of motifs (TAXH, Glu–Thr, and substrate tunnel gating pair) that demonstrate conservation of structural properties of chitin-active LPMOAA10s despite deviations in sequence motifs (Figure 3C). The structure of CbpD reinforces the structurally conserved nature of the Glu residue in the secondary shell that is oriented toward the active site His-brace irrespective of its occurrence in the amino acid sequence, in line with observations for other LPMOAA10s (O’Dell *et al*., 2017; Forsberg *et al*., 2019). While the two residues are flipped compared to the Glu60/Asn185 pair in *Sm*AA10A, the preservation of the pair’s structural position in the AA10 domain indicates Glu174/Asn46 may play a similar role in gating access to the active site when CbpD is bound to crystalline chitin as Glu60/Asn185 do in *SmAA10A* (Bissaro *et al*., 2018*a*). Notably, a previous study found that mutating Glu60 to Asn was deleterious to catalytic activity (Bissaro *et al*., 2020). That study, however, did not examine the effect of a concomitant mutation to Glu of the other residue in the Glu/Asn pair, which likely explains why CbpD retains chitinolytic activity despite having an apparently deleterious Asn residue at position 46 (equivalent to Glu60) and highlights the need for careful spatial as well as sequence considerations when conducting enzymatic mutation studies.

Other features, however, deviate substantially from previously characterized LPMOAA10s (Figure 3C). In the proximity of the active site, the absence of the proposed cluster of three tryptophans important for an ET monooxygenase catalytic mechanism (Forsberg, Røhr *et al*., 2014) and lack of an identified bacterial CDH (Beeson *et al*., 2015) is further evidence that LPMOAA10s do not rely on internal ET or O_2_ but instead use an H_2_O_2_ peroxygenase mechanism to hydrolyze their substrates. These aromatic residues near the active site may still be important, however, in protecting CbpD from oxidative damage (Paradisi *et al*., 2019), and it remains to be seen whether this protective mechanism is why aromatic residues appear to be conserved in general, if not in strict identity, near the active site of LPMOs. The presence of the gating residues similar to *Sm*AA10A (Bissaro *et al*., 2018*a*) indicates that there is a path into the active site of CbpD that is accessible to small molecule reductants, eliminating one concern that originally led to the ET mechanism hypothesis (Li *et al*., 2012; Courtade *et al*., 2016). What remains unclear, however, is whether LPMOs may have reducing agent specificity in addition to substrate specificity (Meier *et al*., 2018; Frommhagen *et al*., 2016). Askarian *et. al*. demonstrated that CbpD chitinolytic activity was enhanced by ascorbate and the redox-active compound pyocyanin, which is secreted by *P. aeruginosa* (Askarian *et al*., 2021). CbpD and other LPMOs may favor small molecule reductants that are secreted by their hosts. Future mechanistic studies can use the structure of CbpD to investigate whether proposed mechanisms are universal and could occur in structurally diverse LPMOs with the same and divergent substrates or whether mechanisms may be family-dependent and whether LPMOs utilize general or context-specific reductants if an H_2_O_2_-based mechanism is employed.

The presence of Arg169 in the active site secondary shell and absence of a pocket on the substrate surface is also noteworthy because previously, an Arg in this position was associated only with cellulose-activity in LPMOAA10s (Forsberg, Røhr *et al*., 2014). Interestingly, an Arg is present in this position in two other chitin-active LPMOAA10s (Tma12 and *Cj*LPMO10A) (Supplementary Figure 2), all three of which are structurally similar to the cellulose active LPMOAA10 *Sc*LPMO10C based on the DALI server “all against all” comparison (Holm, 2020) and are the only members of their structural class to lack cellulolytic activity (Figure S2). This raises the possibility that there are distinct yet convergent features of LPMOAA10s that have evolved to confer chitin-specific activity and indicates these features may have not all been identified. Interestingly, CbpD may have a shallow pocket just to the side of the active site that may be able to accommodate an N-acetyl group (Figure 4C), although it is between the putative channel gating resides (Glu174/Asn46) and the copper active site and may be part of the substrate tunnel when CbpD is bound to chitin. The structures of CbpD, Tma12, and *Cj*LPMP10A now provide three examples of LPMOAA10s that are strictly active on chitin yet lack the pocket previously thought to either facilitate substrate hydrolysis by accommodating the oxygen cosubstrate or substrate binding by accommodating the N-acetyl group present in chitin and should inform future investigations into the deterministic features that confer substrate specificity in LPMOAA10s.

The variation of the chitin-specific L2 motif Y(W)EPQSVE on the substrate binding surface of CbpD further challenges previously strong conclusions about the deterministic features of LPMOAA10s that confer substrate selectivity. It is unclear whether the L2 motif is more tolerant of sequence variation than first supposed or whether the L2 motif plays a critical role in substrate specificity among LPMOAA10s. These discrepancies in the active site secondary coordination sphere and substrate binding surface also raise an additional question about whether CbpD may be able to bind and hydrolyze substrates beyond chitin.

Many of these studies drew their conclusions about discriminatory features from a relatively small number (10s) of evolutionarily closely-related LPMO sequences. The structure of CbpD highlights the importance of including an evolutionarily diverse array of LPMOs in the identification and analyses of substrate specificity motifs. This is particularly true given that the bacterial LPMOAA10 CbpD is structurally most similar to the fern LPMOAA10 Tma12 (Figure S2). The presence of the same L2 motif sequence from CbpD in the structurally similar yet evolutionarily distant fern LPMOAA10 Tma12 demonstrates that the structural features of LPMOAA10s that enable chitin binding and degradation have yet to be fully understood and require further investigation among structurally diverse members of the AA10 family.

Even with the structures of CbpD, Tma12, and *Cj*LPMO10A challenging previously identified substrate-specificity deterministic features, there are still very few LPMOAA10s that have known substrate specificity. To date, the CAZy Database contains over 8000 LPMOAA10 proteins, yet only 32 are listed as “characterized”, and this list does not include CbpD, despite its substrate binding being characterized in 2000 (Folders *et al*., 2000) and its catalytic activity profile confirmed in 2021 (Askarian *et al*., 2021). One characterized LPMOAA10, *Kp*LPMO10A, appears to be able to cleave chitin, cellulose, and xylan (Corrêa *et al*., 2019), expanding the substrate scope of LPMOAA10s beyond just chitin and cellulose, yet no studies have investigated what structural features may enable this broader substrate scope in an LPMOAA10. Until a larger portion of the LPMOAA10 family has characterized substrate profiles, any deterministic motifs and features identified among LPMOAA10s must be taken as indicative, but not deterministic, at best.

Because the catalytic activity of CbpD is necessary for *P. aeruginosa* virulence (Askarian *et al*., 2021), discrepancies in putative substrate specificity features highlight the importance of considering the role of LPMOs in both carbohydrate polymer degradation and virulence when identifying the residues and motifs that confer substrate specificity. Humans do not make the chitin polymer, but do have the highly glycosylated mucin protein. Polymicrobial biofilms in which *P. aeruginosa* interacts with fungi could also be an opportunity for CbpD activity as fungal cell walls are chitinous.

The presence of a number of discrepancies in previously identified substrate-specificity motifs and its role in virulence mean CbpD should serve as a basis for more extensive modeling of LPMOAA10s interacting with chitin. LPMOAA10–chitin modeling to date has only been performed with single-domain LPMOAA10s (Bissaro *et al*., 2018*a*). Additionally, this modeling was performed without the consideration of post-translational modifications (PTMs). The structure of CbpD, however, reveals that two residues that are phosphorylated when CbpD is secreted from *P. aeruginosa* (Ouidir *et al*., 2014)—Ser172 and Tyr40—form part of the chitin-binding surface of CbpD (Figure 4), and in the case of Tyr40, are critical for substrate binding (Beeson *et al*., 2015; Bissaro *et al*., 2018*a*). It has been speculated that phosphorylation at Ser172 proximal to E174 in the secondary coordination sphere of the Cu active site likely affects cosubstrate (O_2_ or H_2_O_2_) activation and may alter the Cu redox potential (Askarian *et al*., 2021). A number of these post-transitionally modified residues are also in the AA10:GbpA2 domain interface of CbpD, likely encouraging a more elongated overall conformation, similar to the observed solution scattering conformation. Additionally, a recent bioinformatics analysis of 27,000 LPMOs found that most of the eight LPMO AA families are enriched for Ser and Thr residues in C-terminal regions that are prime candidates for PTM, specifically O*-*glycosylation (Tamburrini *et al*., 2021), which may affect secretion (Vorkapic *et al*., 2019). Of note, the Tma12 structure has an N-glycosylation at Asn158, structurally equivalent to Ser136 in CbpD, which has been identified as phosphorylated when CbpD is expressed and secreted by *P. aeruginosa* (Ouidir *et al*., 2014). A complete understanding of how LMPOAA10s are secreted by their native organisms and bind their substrates cannot be obtained without taking these PTMs into consideration.

The β-strand interaction observed between GbpA2 domains of crystallographic symmetry mates raises another question about the possible biological function of structural features of CbpD. Edge β-strands are well-known to mediate protein:protein interactions, which in some cases are dynamic and interchangeable (Miyazaki *et al*., 2021; Richardson & Richardson, 2002; Monteiro *et al*., 2013; Yu *et al*., 2014). Given the broad substrate repertoire of the *P. aeruginosa* T2SS (Cianciotto & White, 2017), and the expectation that recognition of secreted proteins by the multiple proteins along the secretion machinery pathway relies on dynamic, short-lived interactions (Douzi *et al*., 2011; Michel-Souzy *et al*., 2018) as well as inherent disorder (Gu *et al*., 2017; Pineau *et al*., 2021), it is interesting to imagine that the β15 edge strand of CbpD may serve as a handle for interaction with secretion machinery proteins along the pathway through the cell envelope. The CbpD structure may thus provide an experimental entrée into a better understanding of how T2SS substrates are recognized.

## 5. Conclusions

In addition to demonstrating the utility of new structural biology tools, the structure of CbpD provides evidence to both support and challenge aspects of the current model for how LPMOAA10s bind their substrates, specifically ones that are preferential for chitin. The absence of the CBM73 domain in the structure also raises questions about the inherent stability of smaller carbohydrate binding domains, and a crystal structure of a CBM73 domain remains to be solved. One possibility is to perform high-resolution, single particle electron microscopy of CbpD bound to chitin fibrils. This could provide the structure of full-length CbpD and shed light on the conformation of CbpD when bound to crystalline chitin. A better understanding of the structure, conformations, and dynamics of multimodular LPMOs could lead to advances in understanding the role of these enzymes in biomass degradation and virulence.

## Acknowledgements

We thank Dr. Silvia Spinelli for expert assistance with data collection and Dr. Brian Fox for careful reading of the manuscript. We acknowledge funding from the ANR to RV and KTF (ANR-14-CE09-0027-01) and to RV and Olivera Francetic (ANR-19-CE11-0020-01), from the UW-Madison Foundation E. B. Fred Professorship and the USDA (Hatch Act Formula Fund WIS03052) to KTF. This project was supported in part by a fellowship award to CMD through the National Defense Science and Engineering Graduate (NDSEG) Fellowship Program, sponsored by the Air Force Research Laboratory (AFRL), the Office of Naval Research (ONR) and the Army Research Office (ARO).

