## Supplementary figures for "CbpD crystal structure adds intrigue to substrate-specificity motifs in chitin-active lytic polysaccharide monooxygenases"

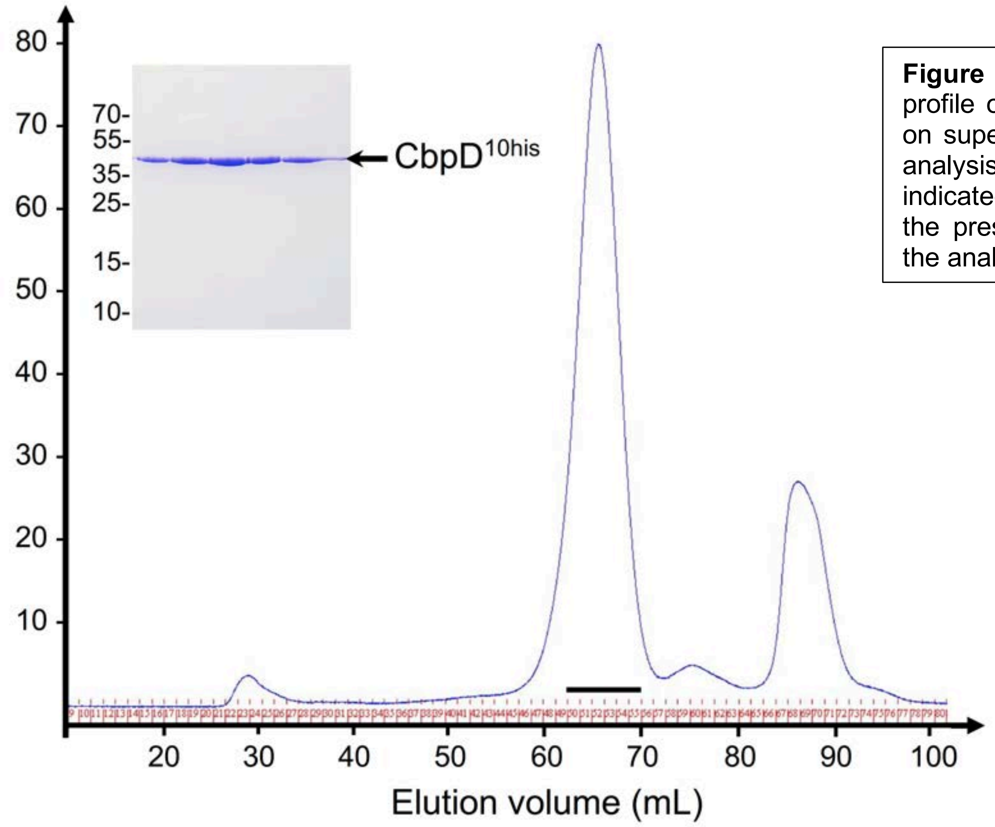


**Figure S1** CbpD purification: SEC profile of CbpDHis purified by IMAC on superose 200 16/60. SDS-PAGE analysis of the SEC fractions indicated by the black line revealed the presence of full-length CbpD in the analyzed peak (43kDa).

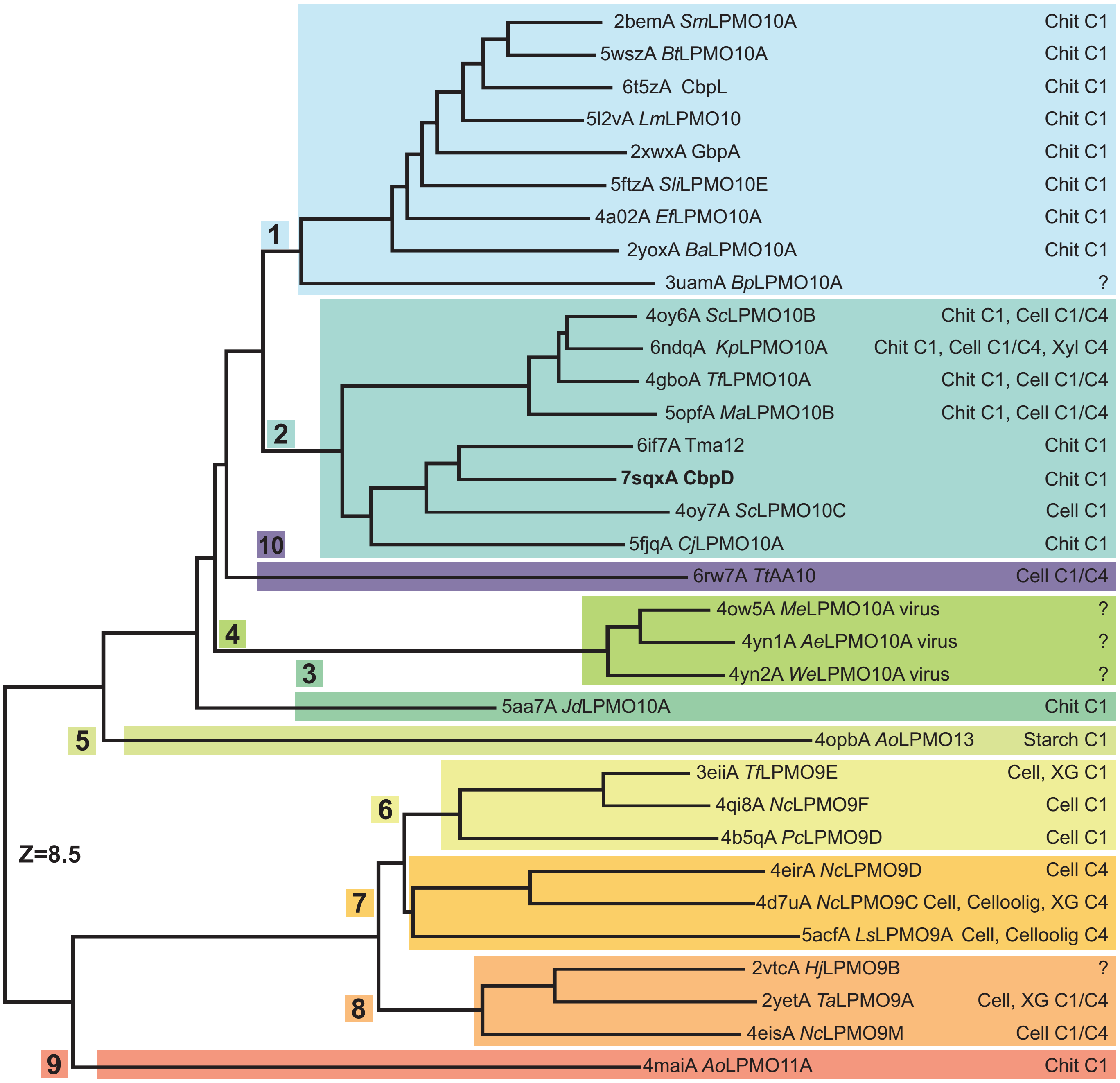

2bemA *Sm*LPMO10A Chit C1
5wszA *Bt*LPMO10A Chit C1
6t5zA CbpL Chit C1
5l2vA *Lm*LPMO10 Chit C1
2xwxA GbpA Chit C1
5ftzA *Sli*LPMO10E Chit C1
4a02A *Ef*LPMO10A Chit C1
2yoxA *Ba*LPMO10A Chit C1
3uamA *Bp*LPMO10A ?
4oy6A *Sc*LPMO10B Chit C1, Cell C1/C4
6ndqA *Kp*LPMO10A Chit C1, Cell C1/C4, Xyl C4
4gboA *Tf*LPMO10A Chit C1, Cell C1/C4
5opfA *Ma*LPMO10B Chit C1, Cell C1/C4
6if7A Tma12 Chit C1
**7sqxA CbpD** Chit C1
4oy7A *Sc*LPMO10C Cell C1
5fjqA *Cj*LPMO10A Chit C1
6rw7A *Tt*AA10 Cell C1/C4
4ow5A *Me*LPMO10A virus ?
4yn1A *Ae*LPMO10A virus ?
4yn2A *We*LPMO10A virus ?
5aa7A *Jd*LPMO10A Chit C1
4opbA *Ao*LPMO13 Starch C1
3eiiA *Tf*LPMO9E Cell, XG C1
4qi8A *Nc*LPMO9F Cell C1
4b5qA *Pc*LPMO9D Cell C1
4eirA *Nc*LPMO9D Cell C4
4d7uA *Nc*LPMO9C Cell, Celloolig, XG C4
5acfA *Ls*LPMO9A Cell, Celloolig C4
2vtcA *Hj*LPMO9B ?
2yetA *Ta*LPMO9A Cell, XG C1/C4
4eisA *Nc*LPMO9M Cell C1/C4
4maiA *Ao*LPMO11A Chit C1
1
2
10
4
3
5
6
7
8
9
Z=8.5

**Figure S2** The structural diversity of LPMOs: The expanded dendrogram of unique LPMO crystal structures from Vaaje-Kolstad *et al.* 2017 in addition to CbpD (bold) and subsequent LMPOAA10 structures determined and listed in the CAZy AA10 database. CbpD is a member of structural cluster 2, and *Tt*AA10 may represent a possible 10th structural cluster. Structures are identified by their PDB identifier and the chain ID, followed by their known substrate(s). The scale indicates the DALI Z-score. Some LPMOs have been referred to by other names in the literature, which are indicated in parenthesis; *Sm*LPMO10A (CBP21), CbpL (*Pl*LPMO10A), GbpA (*Vc*LPMO10B, VcAA10B), *Ef*LPMO10A (EfCBM33A, EfaCBM33), *Ba*LPMO10A (BaAA10A, ChbB, BaCBM33), *Tf*LPMO10A (E7), *Sc*LPMO10C (CelS2, ScAA10C), *Ao*LPMO13 (Ao(AA13)), *Tt*LPMO9E (TtGH61E), *Pc*LPMO9D (PcGH61D), *Nc*LPMO9D (PMO-2, NCU01050), *Nc*LPMO9C (NCU02916), *Ls*LPMO9A (Ls(AA9)A), *Hj*LPMO9B (EG7, Cel61B), *Ta*LPMO9A (TaGH61A), *Nc*LPMO9M (PMO-3, NCU07898), *Ao*LPMO11 (Ao(AA11)). Following the naming convention of subsequently discovered and characterized LPMOs, CbpD may also be referred to as *Pa*LPMO10A, Tma12 may also be named *Tm*LPMO10A, and TtAA10 may be called *Tt*LPMO10A.

The organism of each LPMO is:
*Serratia marcescens* (*Sm*LPMO10A); *Bacillus thuringiensis* Serovar *kurstaki* (*Bt*LPMO10A); *Photorhabdus luminescens* (CbpL)*; Listeria monocytogenes* (*Lm*LPMO10)*; Vibrio cholera* (GpbA)*; Streptomyces lividans* (*Sli*LPMO10E)*; Enterococcus faecalis* (*Ef*AA10A); *Bacillus amyloliquefaciens* (*Ba*AA10A); *Burkholderia pseudomallei* (*Bp*AA10A)*; Streptomyces coelicolor* (*Sc*LPMO10B, *Sc*LPMO10C)*; Kitasatospora papulose* (*Kp*LPMO10A)*; Thermobifida fusca* (*Tf*LPMO10A)*; Micromonospora aurantiaca* (*Ma*LPMO10B)*; Tectaria macrodonta* (Tma12)*; Pseudomonas aeruginosa* (CbpD); *Cellvibrio japonicus* (*Cj*LPMO10A)*; Teredinibacter turnerae* (*Tt*AA10A); unidentified entomopoxvirus (*Me*LPMO10A, *We*LPMO10A)*; Anomala cuprea entomopoxvirus* (*Ae*LPMO10A)*; Jonesia denitrificans* (*Jd*LPMO10A); *Aspergillus oryzae* (*Ao*LPMO13, *Ao*LPMO11A); *Thermothielavioides terrestris* (*Tf*LPMO9E)*; Neurospora crassa* (*Nc*LPMO9F, *Nc*LPMO9D, *Nc*LPMO9C, *Nc*LPMO9M)*; Phanerodontia chrysosporium* (*Pc*LPMO9D)*; Lentinus similis* (*Ls*LPMO9A); *Trichoderma reesei* (*Hj*LPMO9B)*; Thermoascus aurantiacus* (*Ta*LPMO9A)

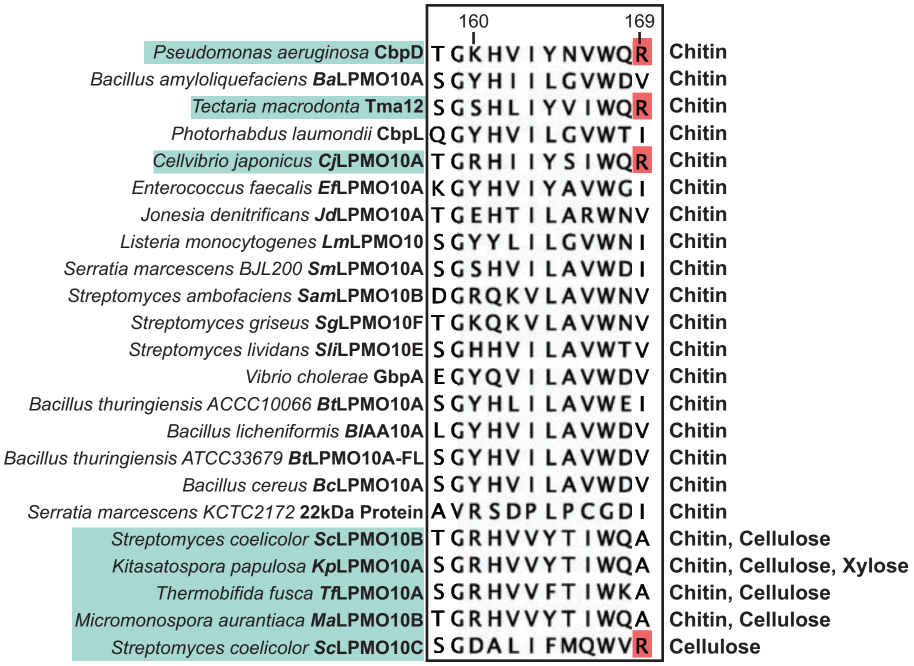


**Figure S3** Position 169 in chitin-active sequences: While previous studies have identified that a short, aliphatic residue is conserved in chitin-active LPMOAA10s (residue 169 in CbpD), creating a shallow cavity on the substrate binding surface that could either accommodate the oxygen species cosubstrate or the N-acetyl group of chitin, the sequence alignment of the chitin-active chitin-active LPMOAA10s listed as characterized in the CAZy database, in addition to CbpD and the other members of the LPMO structural cluster 2 shows Arg (red boxes) is present in 16.7% of chitin-active characterized LPMOAA10s. Position numbering is based on the sequence of CbpD. Boxed LPMO names are members of the LPMO structural cluster 2 (Figure S4). The characterized substrate for each LPMOAA10 is provided to the right of the sequence alignment.

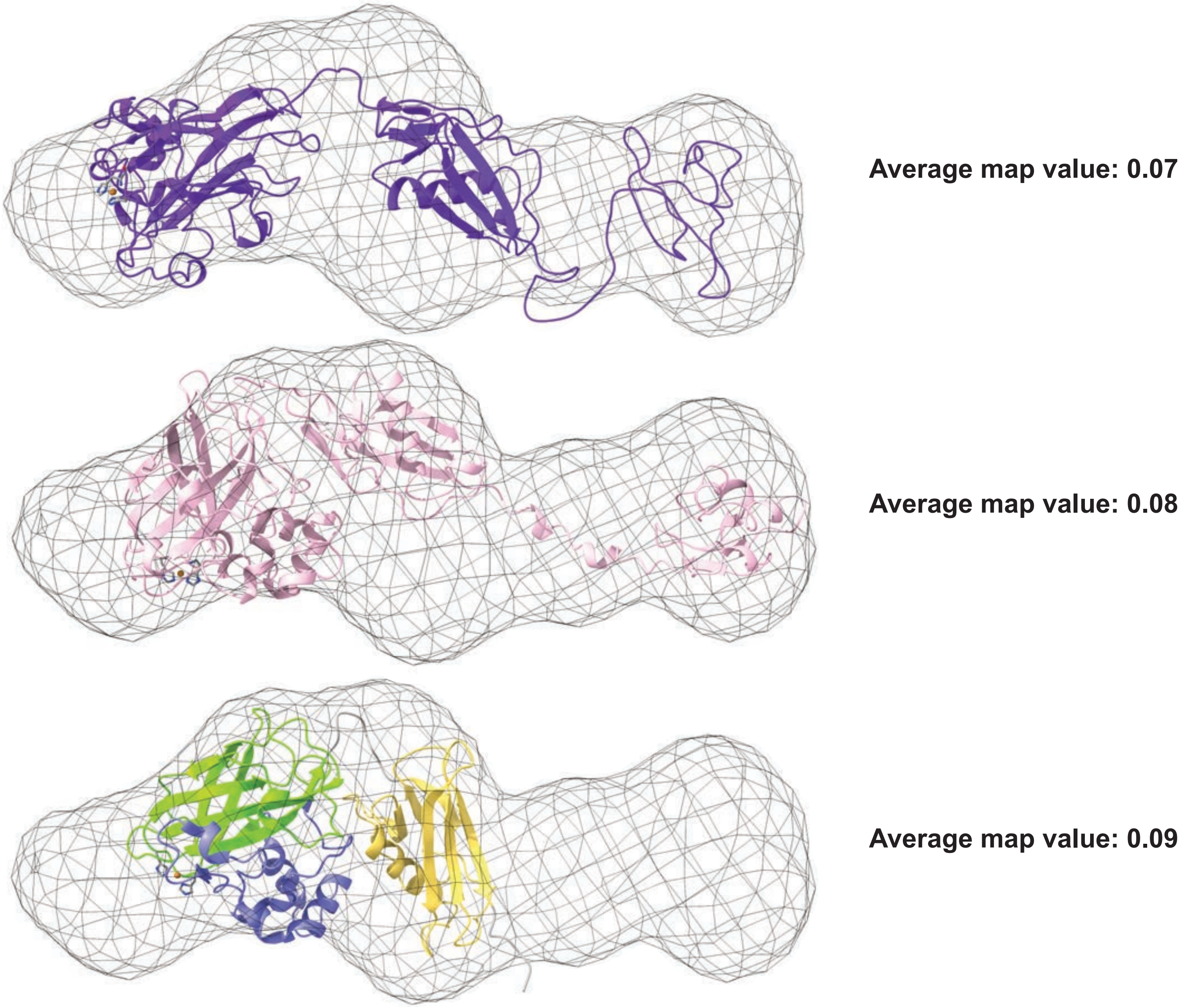


**Figure S4**: The crystal structure of CbpD adopts a more compact conformation than either observed in solution or predicted by AI: Pepsi-SAXS model (Askarian *et al.*, 2021) (SASBDB ID: SASDK42) (top, purple); RoseTTAFold model (middle, pink); and crystal structure (bottom, colored as in Figure2A) of CbpD fit into the 15 Å resolution volume map of the CbpD SAXS model envelope generated by Askarian *et al.* (2021). Average map values after fitting each structure into the density map are provided (right). His brace (sticks) and a Cu ion (sphere) placed in the active site are shown for orientation.
